# bronko: ultrafast, alignment-free detection of viral genome variation

**DOI:** 10.64898/2025.12.01.691650

**Authors:** Ryan D. Doughty, Michael J. Tisza, Todd J. Treangen

## Abstract

As viral sequencing datasets continue to grow, traditional alignment-based variant calling pipelines are becoming computationally prohibitive. To address these challenges, we developed *bronko*, an ultrafast alignment-free framework for detecting viral variation directly from sequencing data. The novel computational approach implemented in *bronko* allows scaling to massive viral sequencing datasets and has three key components: i) a locality-sensitive bucketing function to rapidly identify single-nucleotide polymorphisms (SNPs) relative to reference(s), ii) a direct k-mer count pseudo-mapping approach that approximates a pileup without alignment, and iii) a streaming-based sliding window outlier test to estimate baseline noise across the genome and precisely differentiate real minor variants from noise. Together, these components yield near-linear computational complexity with respect to sequencing depth, enabling *bronko* to process thousands of viral samples rapidly on modest hardware. Our results are threefold: 1) On simulated amplicon sequencing, *bronko* recovers variants with higher precision and comparable recall to existing tools while running up to one to three orders of magnitude faster; 2) *bronko* generates sequence alignments directly from sequencing data, with SNP content similar to that of whole-genome alignment while also running in a fraction of the time, and 3) applying *bronko* to longitudinal sequencing data from chronically infected SARS-CoV-2 patients revealed consistent patterns of intrahost diversification and adaptive mutations over time. Altogether, these results demonstrate *bronko*’s potential as a scalable tool for large-scale viral genomic analyses, overcoming longstanding computational barriers for intrahost and interhost characterization of viral variation.

**Availability:** *bronko* is implemented in Rust and publicly available at https://github.com/treangenlab/bronko or via conda at https://anaconda.org/channels/bioconda/packages/bronko/overview. All results, evaluations, and other code used in this study are available at https://github.com/treangenlab/bronko-test.

## 1 Introduction

Viral sequencing is now central to modern epidemiology, providing unprecedented insight into viral evolution, transmission, and adaptation [1]. By characterizing viral genetic variation across populations, sequencing enables detection of emerging variants of concern, inference of transmission chains, and assessment of selective pressures [2]. These capabilities proved essential during the COVID-19 pandemic, where global monitoring of SARS-CoV-2 variants directly guided vaccine updates, public health interventions, and our understanding of viral dynamics [3]. Beyond major circulating variants, patient-level sequencing data also encodes information on intrahost variants (iSNVs), which can provide additional insight into viral evolution [4]. For instance, iSNVs have been used to reveal early mutational events that can later fix and propagate through populations as well as identify individual transmission events [5–7]. As sequencing continues to become faster and cheaper, the opportunity to continuously monitor viral evolution worldwide in real-time has never been greater.

Standard variant calling pipelines depend on read alignment, followed by identification of both consensus (major) and intrahost (minor) variants. Even with recent optimized algorithms and software [8–11], both read alignment and variant calling remain computationally expensive affairs in genomics workflows [12–14]. Given the NCBI Sequence Read Archive already hosts over 7 million SARS-CoV-2 sequencing datasets alone (∽500 terabytes of data), the current alignment-based paradigm is impractical for population-scale studies of viral variation. This motivates the development of methods that can analyze viral sequencing data more efficiently with similar sensitivity and precision.

Over the years, several alignment-free variant calling methods have been developed for human and bacterial genomics. Rather than aligning reads and calling variants, these approaches detect variation directly from reads, either within or across samples, typically trading a modest loss in sensitivity for substantial runtime gains. Many of these tools focus on discovering novel variation that alignment-based methods may miss, using graph-based data structures to capture local sequence differences [15–20]. Other methods take explicit k-mer–centric approaches to detect smaller variants. Kestrel [21] detects small variants by comparing k-mer abundance profiles between a sample and a reference, enabling rapid inference of sequence differences from coverage imbalances. LAVA [13] clusters k-mers with small edit distances to detect known SNP targets directly. Similarly, kSNP [22, 23] and SKA2 [24, 25] both rely on split k-mers centered on single variable positions, allowing identification of a mutation when the context around the variant remains unchanged.

While these alignment-free methods have shown promise in human and bacterial genomics, viral genomes present additional unique challenges. First, viral sequencing datasets can often reach ultra-high depths (>10,000x), amplifying computational inefficiencies in both read alignment and downstream variant calling [12]. More critically, however, most existing alignment-free methods focus on consensus-level variation and overlook low-frequency iSNVs that are present in most viral sequencing datasets. For viral samples, where some true minor allele frequencies may be below 1%, distinguishing genuine biological signal from sequencing noise and other artifacts requires explicit modeling of error distributions and coverage heterogeneity, particularly in an alignment-free framework where positional context can be limited [26]. Although numerous low frequency variant calling approaches have been developed to handle this task, none have been employed in an alignment-free context [27–29]. Fortunately, viral genome analysis provides numerous opportunities for computational optimization as well. First, their compact genome size enables efficient in-memory representations and exhaustive k-mer indexing approaches. Similarly, despite their rapid mutation rate, the mutational spectrum of most RNA and DNA viruses is dominated by single-nucleotide substitutions, with relatively few structural variants or long insertions or deletions [30]. These simplicities make viral genomes particularly well-suited to k-mer–centric approaches that can capture local nucleotide changes directly from sequencing reads.

Here we introduce *bronko*, a scalable, alignment-free framework to detect both major and minor single nucleotide variants directly from viral sequencing data. *bronko* uses a locality-sensitive bucketing (LSB) function to rapidly identify k-mers within one edit distance of each other [31]. The use of this function enables *bronko* to perform rapid pseudo-mapping of k-mer counts from viral sequencing data directly to an approximate pileup of each genomic coordinate for one or more reference genomes, bypassing read alignment and SAM manipulation entirely. *bronko* then employs a streaming outlier-based statistical model to distinguish genuine minor variants from sequencing noise, adapting across the genome to local error, mutational profiles, and coverage fluctuations. Altogether, these design choices enable *bronko* to run ultra-efficiently and scale near linearly with sequencing depth, making it well-suited for large-scale viral genomic surveillance, longitudinal infection studies, and rapid screening of emerging variants in environmental or clinical contexts.

## 2 Methods

Our method *bronko* performs two primary tasks: (1) rapid pseudo-mapping of k-mer counts to reference genomes of interest, and (2) precise detection of major single-nucleotide variants (SNVs) and intra-host SNVs (iSNVs) from the pseudo-mapping output. Pseudo-mapping is comprised of three components: (i) a (1, 2)-sensitive LSB function that groups near-identical k-mers under fixed edit-distance constraints, enabling single-base mismatch detection (Fig. 1A); (ii) a bucket-based index that stores positional information of all potential variation across references (Fig. 1B); and (iii) a single-pass pileup construction algorithm that converts observed k-mer counts into per-base coverages without explicit read alignment (Fig. 1C). Variant detection operates on the constructed pileups and also has three main stages: (i) reference selection by sequence identity; (ii) estimation of a baseline sequencing-error model using an outlier-based statistic computed in a sliding window across the genome; and (iii) variant calling that combines the baseline noise model, strand-bias metrics, and k-mer support at each position to report SNVs/iSNVs. When provided multiple samples, *bronko* also has the additional ability to generate a multiple sequence alignment from the intersection of called SNVs, enabling rapid large-scale inference of approximate phylogenetic trees.

**Fig. 1.**
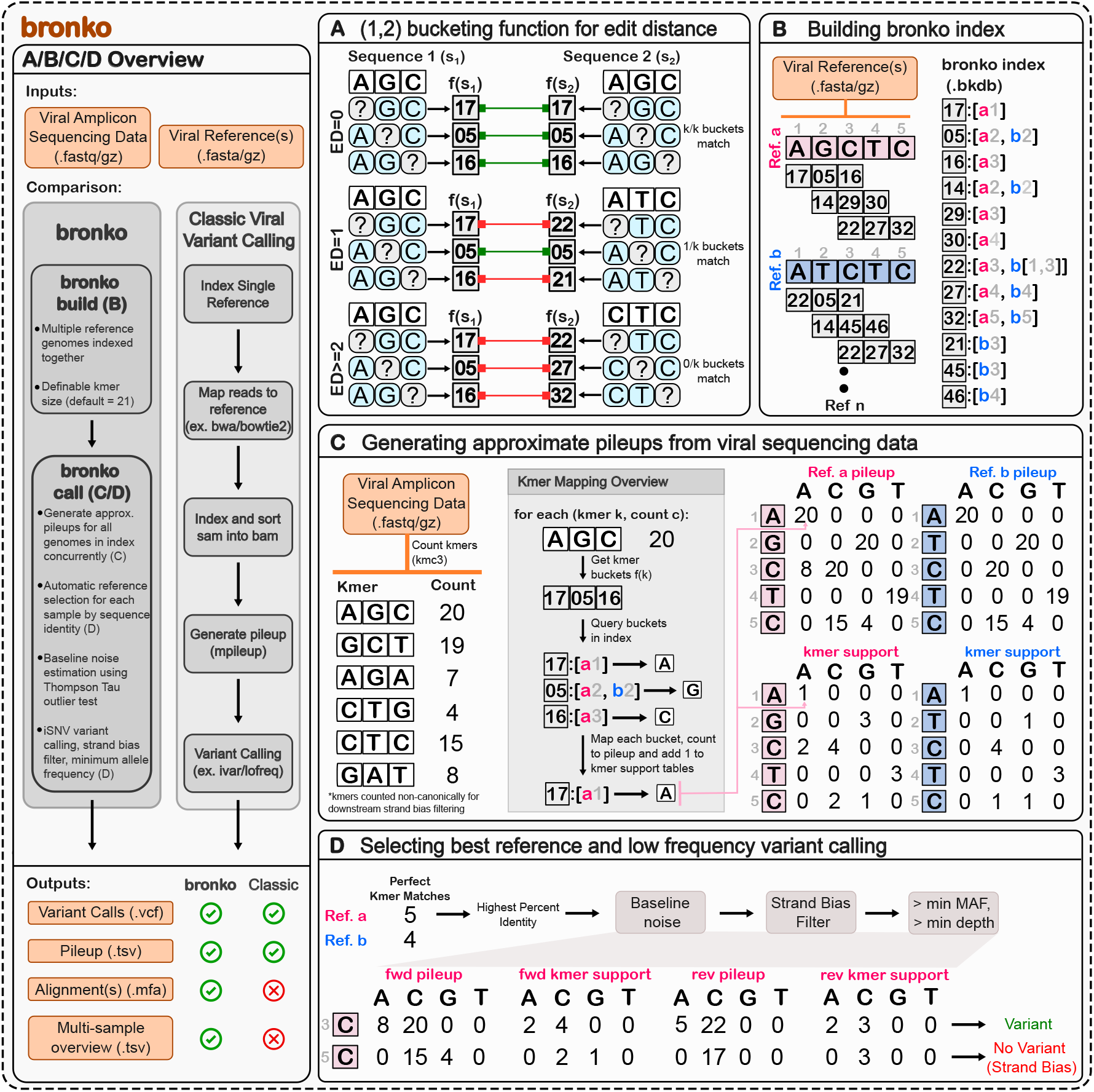
Overview of bronko **a**. (1,2) LSB function *f* for the edit distance. Three different edit distances are shown. White boxes represent k-mers, blue and gray ovals represent variants, and gray boxes represent the bucket values that are returned from f. **b**. Building a *bronko* database from a set of references. Every k-mer in each reference is transformed into buckets, before being placed in a hash table with the reference(s) and position(s) **c**. Generating approximate pileups and k-mer support tables from k-mer counts. Buckets are retrieved for each k-mer before being queried in the index. These positions are then mapped into the pileup and k-mer support **d** Reference selection and variant calling. The reference with the highest identity to each sample is selected before baseline noise, strand bias, and minimum AF/depth filters to identify SNPs and iSNVs

### 2.1 (1,2) Locality-Sensitive Bucketing Functions for Edit Distance

To sensitively detect single-nucleotide variation (SNVs) within viral sequencing data, we utilize a (1,2) LSB function, denoted *f* (*s*), following the formulation described in [31]. In contrast to traditional k-mer matching approaches that commonly rely on exact matches, the LSB approach allows us to efficiently group near-identical k-mers into common “buckets”, while maintaining fixed edit-distance constraints. In *bronko*, we utilize a (1, 2)-sensitive bucketing scheme, which guarantees that any pair of k-mers with edit distance ≤ 1 map to at least one common bucket, while any pair differing by two or more edits (≥2) are assigned to disjoint bucket sets. Formally, for each k-mer *s* of length *k*, the function *f* (*s*) computes a ordered set of *k* integer-valued bucket identifiers *B*_*s*_ = (*b*_0_, *b*_1_, …, *b*_*k*−1_) where each *b*_*i*_ ∈ [0, *k* × 4^*k*−1^]. Intuitively, each bucket *b*_*i*_ is a unique identifier that represents if position *i* in the k-mer was a variant position, and the rest of the k-mer remained the same (see **Figure 1A**). Thus, this formulation allows for identification of not only exact matches between k-mers, but also single nucleotide variation.

For two k-mers *k*_1_ and *k*_2_ with corresponding bucket sets *B*_1_ and *B*_2_, the size of intersection *B*_1_ ∩*B*_2_ points directly to the edit distance of the two sequences. Additionally, in the case of a single mismatch (ED = 1), the matching bucket provides the precise position of the differing base. Altogether, the following 3 cases can be defined for two k-mers, where *ED*(*k*_1_, *k*_2_) represents the edit distance between the two k-mers:

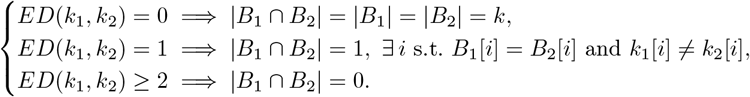

This framework enables identification of single nucleotide variation directly from k-mers in *O*(*k*) time, which altogether allows *bronko* to rapidly identify most single-nucleotide variants directly from k-mers rather than aligning reads.

By default, we adopt a k-mer length of *k* = 21, although this parameter is tunable. Empirically, we found *k* = 21 to provide the best balance between reducing intra-genomic and inter-genomic collisions while also retaining adequate resolution for variant detection. This choice imposes a strict sensitivity limit in that the central variant(s) will be missed in k-length regions with ≥ 3 variants. To ensure intra-genomic collisions do not affect downstream variant calling, we exclude any buckets that occur more than once within a given reference genome, ensuring that each retained bucket corresponds to a unique sequence location. Because we can use the remainder of the k-mer context to identify variants, this has a negligible impact as long as the k-mer size is large enough such that the intra-genomic repeats are rare.

### 2.2 Indexing viral reference sequences

In order to enable rapid k-mer pseudo-mapping to a reference efficiently, we begin by building an index on the buckets of a set of references (see **Figure 1B**). Formally, given a set of *n* viral reference sequences *G* = {*G*_1_, *G*_2_, …}, *G*_*n*_, each k-mer in every reference is first transformed into its canonical form before being mapped to *k* buckets using the bucketing function *f*. As each bucket corresponds to a specific nucleotide position within the k-mer, we store the buckets as keys and their associated reference identifiers and positions as values. In practice, we also record whether the k-mer was converted to its reverse complement during canonicalization, allowing us to recover the original strand orientation after matching. While conceptually simple, this indexing scheme enables ultra-rapid identification of most single-nucleotide variants relative to a larger reference set, a capability not included in most existing variant calling pipelines.

### 2.3 Direct k-mer Pseudo-Mapping and Approximate Pileup Construction

Given an index over a collection of viral reference genomes *G* as described above, we directly map observed k-mer counts from amplicon sequencing reads to per-base pileups for all references simultaneously (see **Figure 1C**). We label this process k-mer pseudo-mapping. Rather than performing conventional read alignment followed by a separate variant-calling stage, pseudo-mapping constructs pileups in a single pass by mapping k-mer counts using indexed bucket positions. This eliminates the need for full read alignment while retaining single-nucleotide resolution in most cases. Because deep viral sequencing datasets are often highly redundant, this approach compresses the relevant variation into a single call, avoiding depth-dependent pileups that scale poorly with coverage. Intuitively, mapping k-mers rather than full reads provides a rapid, more linear-scaling form of compression.

For each reference genome *g* ∈ *G* of length *ℓ*_*g*_, we allocate tables to store estimated base coverage and the number of unique k-mers mapping to each position-nucleotide pair (henceforth referred to as k-mer support). We do this for both the forward and reverse strands, resulting in 4 tables. Specifically, we define:

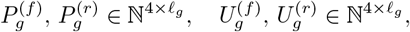

where 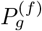 and 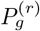 represent the cumulative pileup counts for the forward (f) and reverse (r) orientations, respectively, and 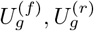 is the k-mer support. All entries are initialized to zero:

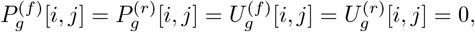

for *i* ∈ {*A, C, G, T* } and 1 ≤ *j* ≤ *ℓ*_*g*_.

In order to map sequencing data to each table, we first enumerate and count all observed k-mers non-canonically from the viral sequencing data using kmc3 [32], yielding a set 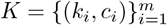 where *k*_*i*_ denotes a unique k-mer and *c*_*i*_ its observed count. We start by discarding k-mers with counts below a small threshold *n*_min_ (default *n*_min_ = 3). Because in most sequencing datasets, a large percentage of k-mers are rare, this filtering step effectively helps to remove potentially spurious k-mers that could otherwise contribute false minor alleles while also speeding up the process by reducing the number of k-mers that need to be mapped.

For each retained k-mer *k* with count *c*_*k*_, we compute its canonical form *k*_*c*_ and subsequently apply the bucketing function *f* (*k*_*c*_) to obtain the ordered set (*b*_0_, *b*_1_, …, *b*_*k*−1_), where each *b*_*i*_ corresponds to the bucket representing a variant position at position *i* of the k-mer. Each bucket *b*_*i*_ is then queried against the index to identify a potential match to the reference at location *j*, along with the strandedness of the match *ρ*. For each such match, we identify the corresponding nucleotide *k*[*i*] in the k-mer and update the appropriate pileup and k-mer support tables:

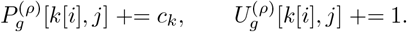

### 2.4 Selecting the best reference for a sample

In the case where the index *G* contains more than a single reference, we automatically select the best reference after pseudo-mapping to each one individually. Given that viral genomes are relatively small and most of the genomes encode proteins, we use a simple heuristic that estimates the percent identity across the full genome of a sample to each reference *g* ∈ *G*. To do this, we calculate the following:

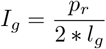

where *p*_*g*_ is the number of k-mers that were mapped perfectly to genome *r* and *l*_*g*_ is the length of the genome. The number of perfect k-mers can be efficiently calculated during the mapping process by counting the k-mers in which all buckets have consecutive positions in the reference. The total is divided by 2 to account for k-mers mapping to both the forward and reverse strands, given k-mers are counted non-canonically. This process ensures that we select a reference that is closest to the actual sample, reducing the number of major variants that could disrupt minor variant calling in the sample.

### 2.5 Low frequency variant calling

Once a reference genome is selected, variant calling can be performed on the pileups for that reference. There have been many methods over the years that use different statistical methods in order to differentiate real mutational signal from sequencing error and other artifacts. We utilize a streaming version of the Thomp-son–Tau outlier test, adapted from the approach implemented in Outlyzer [33]. We selected this method due to its strong performance in external benchmarks [14], showing (i) the highest recall at higher minor allele frequencies (>1%), (ii) high precision at lower minor allele frequencies (<1%), and (iii) good computational efficiency and robustness to changes in sequencing depth. Our implementation extends this approach to a streaming context, allowing continuous evaluation of local noise distributions without recomputing statistics for each sliding window across the genome (see **Figure 2**).

**Fig. 2.**
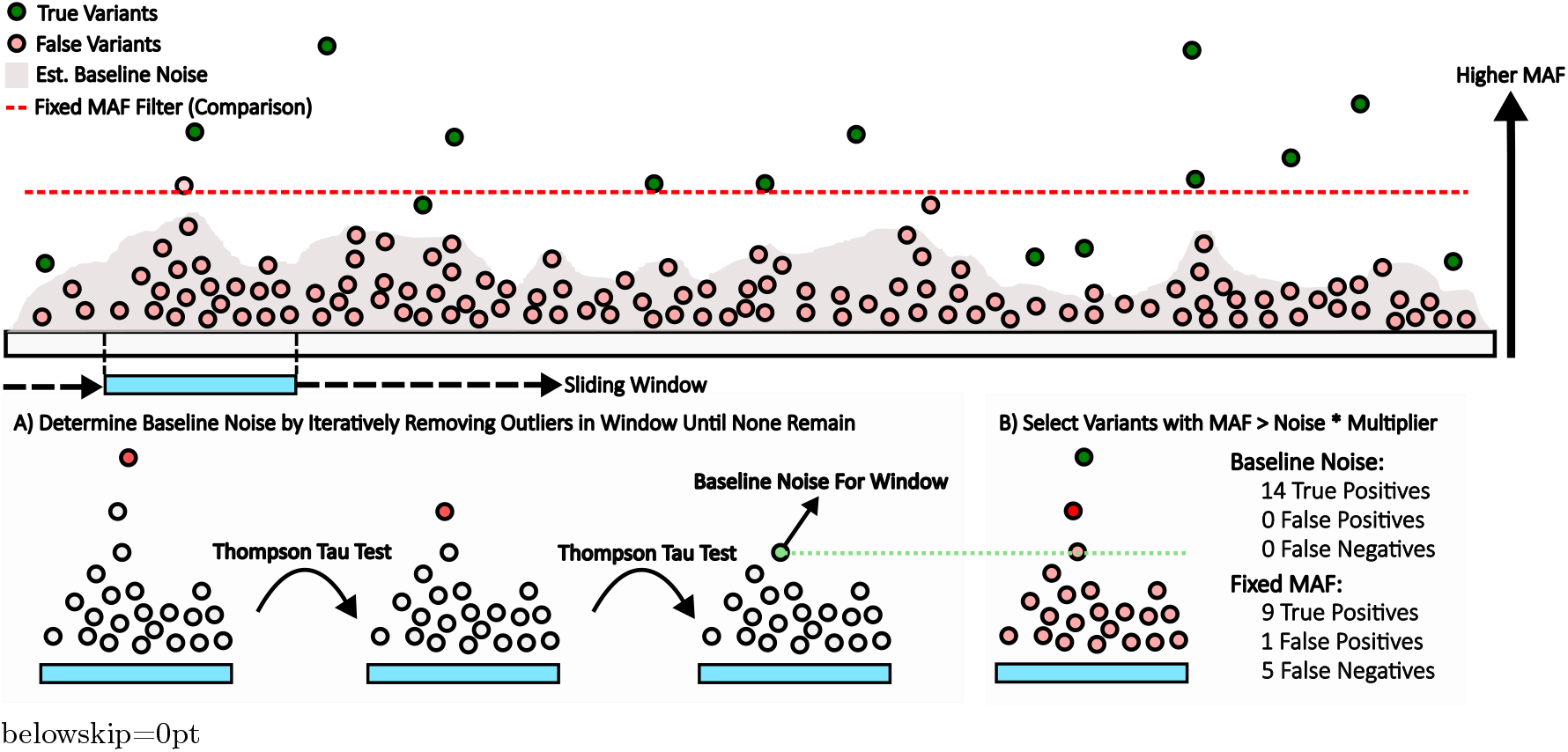
Baseline noise filter for low-frequency variants. Real variants are shown in green, errors in pink. The gray shadow represents the estimated baseline noise, and the red dashed line shows the fixed MAF threshold for comparison. **a**. Sliding window outliers are iteratively removed using a modified Thompson Tau test to estimate local noise **b**. Variants with MAF above the window’s baseline (scaled by a multiplier) are kept, others discarded

Within a sliding window of fixed length *w*, the algorithm maintains running estimates of the number of non-major alleles (*n*_*w*_) with minor allele frequency (MAF) greater than 0, the sum of MAFs (*s*_*w*_), and the sum of squared MAFs 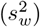. Because there can be at most three minor alleles at each genomic position, all MAFs within the window are stored in an array of length 3*w*. The three possible minor alleles for a given reference position *p* are indexed as *i* = (*p* mod *w*) × 3.

After updating the minor allele frequencies at position *p*, as well as *n*_*w*_, *s*_*w*_, and 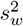, the running mean (*µ*_*w*_) and variance 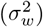 are computed using Welford’s approach for updating variance [34, 35]:

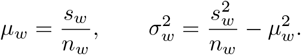

To efficiently remove outliers, two auxiliary data structures are maintained. First, a short, sorted list of length *ℓ* ≪ *w* (default 20) containing the top-*ℓ* MAFs in the current window, and second, a binary indicator array of length 3*w* marking whether each MAF belongs to the top-*ℓ* list. When a new minor allele enters the window, it is compared to the smallest element in the top-*ℓ* list. If larger, it is inserted in sorted order and flagged in the indicator array. Otherwise, it is ignored. When an element exits the window, the indicator is checked, and if present in the top-*ℓ* list, it is removed and the list is adjusted. This structure allows rapid identification of potential outliers without recomputing the max from the entire window.

For each window, baseline sequencing noise is determined by iteratively removing outliers using a modified Thompson–Tau test until no outliers remain, updating the window statistics after each removal. The maximum MAF in the window *x*_*i*_ is removed if *δ*_*i*_ *> τ* ∗ *σ*_*w*_, where *δ*_*i*_ = |*x*_*i*_ − *µ*_*w*_| and the dynamic threshold *τ* is derived from the Student’s *t*-distribution with *n*_*w*_ − 2 degrees of freedom:

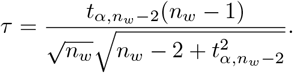

Here, *α* is the desired false-positive rate (we utilize 10^−3^). Because we test for upper-tail outliers only, the one-tailed form of *t*_*α,N*−2_ is used. After each removal, the statistics *n*_*w*_, *s*_*w*_, 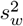, *µ*_*w*_, 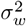, and *τ* are updated, and the next largest MAF is tested. This process continues until the maximum MAF value within the window does not reach the criteria to be considered an outlier.

Once the baseline noise profile is estimated across the genome, candidate variants are retained only if their MAF exceeded the baseline at their position by a specified multiplier. While outlyzer applies a uniform 2x multiplier threshold [33], we found that using a lower default multiplier of 1.5 × and gradually increasing it up to 2 × for low-frequency variants (MAF *<* 1–2%) improved both precision and recall. Formally, the adaptive multiplier for a variant with MAF v is defined as *m*(*v*) = *b*_0_ + *p*_0_*a*^*v*^, where *a* = 0.03 and *p*_0_ = 0.5. This function yields a maximum multiplier of 2 × for MAFs approaching 0 and converges toward 1.5 × for variants near MAF *>* 1.5%.

#### Additional filters for minor variants

In addition to our baseline noise filter, we also employ a strand filter to remove variants with significant strand bias. We utilize the strand odds ratio filter from GATK, although keep a relaxed threshold of <= 6 following guidance from [36]. The GATK strand odds filter was chosen as opposed to other strand frequency filters due to its performance on higher depth sequencing data. We also utilize the number of k-mers supporting any given variant from the tables 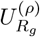. The motivation behind this filter was to remove variants that can pileup from artifacts that near the ends of amplicon sequences. Because we are unable to easily utilize quality information directly from sequencing data, which often would remove variants of this type, we utilize the robustness of multiple k-mers supporting a single position to provide a secondary proxy. By default we require both strands to be present in 2 or more k-mers, although these parameters are configurable.

### 2.6 Multiple Sequence Alignment, Software Outputs and Configuration

When multiple sequencing samples are provided, *bronko* can generate an alignment for each set of sequences aligned to the same reference. To do so, we perform variant calling as described above on each sample independently. If querying against a larger reference set, any genome with > 3 samples mapped to it is deemed suitable for alignment. *bronko* then takes the union of SNV positions (positions with variants > 50% MAF) across all samples for each reference. The consensus variant is taken for each genome at all of these positions, which are then used to build a multiple sequence alignment for that reference.

#### Mapping statistics

For each sample, we also generate some oveview statistics that can assist with quality control in downstream pipelines. The first are the breadth and depth of coverage of each sample. Breadth of coverage is defined as the percentage of bases in the selected reference with a total depth >= 0. This is normalized by the genome length to get a final value from 0 to 1. The depth of coverage reported is simply the average depth across all of those non-zero positions. We also report the number of k-mers that were mapped as “perfect” (edit distance = 0) or “variant” (edit distance = 1) as well as unmapped (edit distance >= 2). These can help provide insight into the quality of the sequencing dataset, although we have found that the number of variant and unmapped k-mers scales well with sequencing depth in practice.

#### Software development and parameters

*bronko* is implemented in Rust and available publicly on bioconda. All experiments in this paper were run using *bronko* v0.1.3 on an AMD EPYC 7742 64-Core Processor using 10 threads unless otherwise noted.

## 3 Results

### 3.1 Low frequency variant calling on simulated amplicon sequencing data

We first assessed *bronko*’s performance in detecting iSNVs using simulated viral sequencing data with a defined ground truth. Ten Human Papillomavirus 16 (HPV16) datasets were generated with GENOMICON-Seq [37], each consisting of one million reads simulated under parameters described in the Appendix. We benchmarked *bronko* against two widely adopted viral variant calling tools: LoFreq [27], and iVar [38]. LoFreq uses base-quality–aware statistical modeling for low-frequency variants, while iVar applies minimal filtering and reports variants above a user-defined MAF threshold. Performance of each tool was evaluated for precision, recall, and F1 score, considering single-nucleotide variants only.

As shown in **Figure 3A**, *bronko* consistenly demonstrated superior precision across a range of increasing MAF thresholds. At MAF ≥ 0.5%, *bronko* achieved perfect precision, and maintained 88% precision even at a 0.1% MAF cutoff. LoFreq showed comparable performance at higher frequencies, also achieving perfect precision above 1% MAF, but dropped slightly below *bronko* with 91% and 76% at 0.05% and 0.01% MAF thresholds respectively. iVar, on the other hand, exhibited lower precision across all three thresholds, with a maximum precision of 43%. In terms of recall, iVar outperformed both *bronko* and LoFreq, primarily due to a high proportion of variants containing MAFs near the error rate of sequencing (0.1%). *bronko* identified slightly more true positives than LoFreq across the simulated HPV datasets, identifying 60 TP variants versus 55 for LoFreq, although the recall is relatively consistent for both tools. iVar was more sensitive than both, identifying 236 true positive variants, reflecting a near 300% increase in TP variants. However, iVar called 1549 false positive variants across the 10 samples in the process. *bronko* had the best or co-best F1 score across all tools and MAF thresholds.

**Fig. 3.**
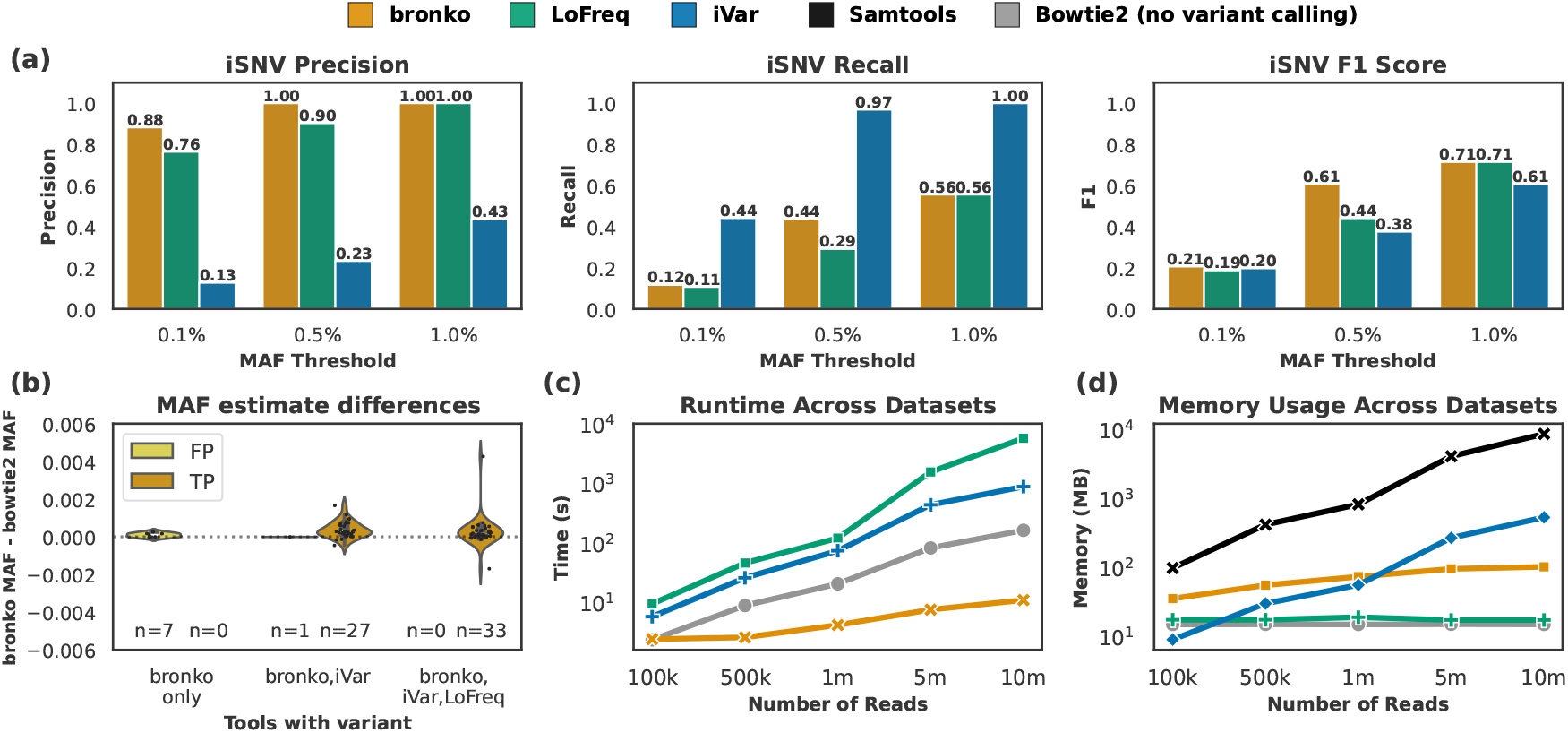
Evaluation of iSNV variant calling. **a**. iSNV precision, recall, and F1 score for *bronko*, LoFreq, and iVar over increasing MAF thresholds **b**. Absolute difference between *bronko* estimated MAF and Bowtie2 MAF for variants called by *bronko*. Calls grouped by tools calling variant and TP/FP **c**. Runtime (seconds) across datasets of increasing size for each tool. Bowtie2 and Samtools time is incorporated into the full time for LoFreq and iVar. **d**. Memory usage across tools. iVar and LoFreq utilize Samtools (black). Memory utilization without this step is shown in displayed colors

Because *bronko* constructs its pileup through the direct pseudo-mapping of k-mer counts rather than traditional alignment, we next evaluated its ability to accurately estimate minor allele frequencies relative to alignment-based methods (see **Figure 3B**). Specifically, we compared the difference between MAFs reported by *bronko* and those derived from Bowtie2-based pileups then used by iVar and LoFreq. Overall, *bronko* exhibited reasonable concordance with Bowtie2, with all minor variants having an absolute discordance of less than 0.005, and 55 of 68 variants (80.4%) falling within 15% difference from the reference MAF. The observed deviations were slightly above the truth on average, suggesting *bronko* introduces a slight positive bias. The distributions were also not different depending on the other tools calling a given variant.

Critical for large scale genomic analyses, *bronko* demonstrated substantial computational advantages over both approaches (see **Figure 3C/D**). In benchmarks with 1 million reads, *bronko* completed variant calling in less than 3 seconds. In comparison, iVar and LoFreq required 73 and 119 seconds respectively, under identical conditions. When the number of reads increases up to 10 million, *bronko* maintains a runtime under 10 seconds, while iVar’s runtime jumps to 890 seconds (14.8 minutes) and LoFreq jumps to 5409 seconds (90.14 minutes), representing nearly a three orders of magnitude speed improvement for *bronko*. In terms of memory utilization, *bronko* consistently utilized under 100 MB, even as the size of the dataset increased. In contrast, both alignment-based pipelines exhibited considerably higher memory consumption, primarily due to SAM/BAM file manipulation using Samtools [39]. At 10 million reads, peak memory for both tools reached 8.5 GB, primarily driven by intermediate SAM file handling. Even when excluding this overhead, iVar still used slightly more memory on larger datasets, primarily because it pipes Samtools mpileup directly into the command. On the other hand, LoFreq alone consistently used under 20 MB regardless of dataset size. However, in the end, both alignment-based pipelines inherently require an intermediate SAM manipulation step, and thus, *bronko* outperforms both methods when taking this into account (**Fig. 3D** black line). As this analysis used a single reference, we also evaluated the performance of the *bronko* build and call commands across different genome lengths and numbers of genomes **(see Supplementary Figs. 1-2**). In short, we found that bronko’s performance only starts to degrade beyond genomes of 1Mbp, and performance scales linearly with the number of reference genomes when the lengths of input genomes are uniform.

**Table 1.** Alignment SNP overlap and performance comparison across simulated datasets.

| Organism<br>(#) | Total Data<br>Size (TB) | Runtime, Memory |  | SNPs |  |  | Shared<br>SNPs (%) |
| --- | --- | --- | --- | --- | --- | --- | --- |
|  |  | Alignment | Bronko | Alignment | Bronko | Shared |  |
| CoV<br>(n=543) | 1.03 | 48.54 hrs<br>2.12 GB | **1.59 hrs**<br>**1.987 GB** | 0 | 0 | 3647 | 100 |
| HIV (S1)<br>(n=41) | 0.012 | 15.88 min<br>**0.35 GB** | **1.99 min**<br>0.74 GB | 1175 | 11 | 9041 | 88.4 |
| HIV (S2)<br>(n=39) | 0.011 | 13.05 min<br>**0.35 GB** | **1.65 min**<br>0.72 GB | 134 | 0 | 3754 | 96.5 |
| HIV (S3)<br>(n=37) | 0.011 | 12.92 min<br>**0.34 GB** | **1.57 min**<br>0.72 GB | 0 | 0 | 664 | 100 |

### 3.2 Evaluation of SNPs from multiple sequence alignments across diverse viral datasets

To evaluate *bronko*’s ability to produce a multiple sequence alignment from multiple viral sequencing datasets, we simulated viral outbreaks and compared the SNPs included in the multiple sequence alignments produced by *bronko* to those included from the core-genome alignment tool Parsnp2 [40]. First, we simulated 552 virus genome sequences emulating the early spread of SARS-CoV-2 (CoV). To do so, we used favites-lite [41], which returns mutated sequences based on a known transmission network. Simulated reads were then generated for each sequence using ART [42] at 30,000x coverage, totaling over 1 TB of sequencing data across all samples. We ran *bronko* to call variants and generate an alignment across all samples with the reference sequence from the origin of the outbreak. Separately, each set of reads was mapped to the reference using Bowtie2 [10] and consensus sequences were generated using Samtools mpileup [39]. Parsnp2 was then used to align the consensus sequences to get a final core-genome alignment for comparison to *bronko*.

*bronko* successfully processed the full dataset in just over 90 minutes, averaging just under 10 seconds per sample. In contrast, the read alignment-based approach required more than 48 hours on 32 cores, with Parsnp2 adding fewer than 3 minutes to produce the final core-genome alignment. Both pipelines used approximately 2 GB of RAM. Notably, the comparison workflow did not include the detection of minor iSNVs, which *bronko* identifies natively and thus is included in the runtime. The SNPs identified by *bronko* and Parsnp2 were identical (3,647 variants total), highlighting strong performance of *bronko* for another use case. However, this dataset, where genomes differed between 2-14 (median 6) SNPs from the reference, represents a relatively simple case for *bronko*’s algorithm as long as there are no local hot spots of variation. To evaluate our approach on a more challenging dataset, we repeated this experiment with three smaller sets of simulated HIV genomes (full details and discussion available in supplementary materials). In short, we simulated three datasets (S1, S2, and S3), varying only the height parameter in the Non-Homogeneous Yule model used to generate the HIV phylogeny (36, 12, and 2, respectively). This design allowed us to assess how increasing divergence between reference and samples affects *bronko*’s performance, with the three sets representing 3%, 1%, and 0.2% divergence, respectively. Overall, we found *bronko* to perform better as the divergence decreased, with only a mild drop in sensitivity for increased divergence. A large percentage of the false negatives could be directly attributed to decreased sensitivity within k-length regions containing ≥ 3 variants (**see Supplementary Table 1 and Supplementary 1.3**).

### 3.3 Tracking viral evolution within chronically-infected SARS-CoV-2 patients

To demonstrate a real-world application of *bronko*, we re-analyzed a subset of a dataset from a recent study of chronically infected SARS-CoV-2 patients from the UK [43]. In the original work, the authors primarily investigated variants with a MAF ≥ 20%, citing low viral titers in many samples. We focused on high-quality samples (mean sequencing depth ≥ 3000× and breadth of coverage ≥ 95%) to investigate potentially relevant low-frequency variants (1–20% MAF). We started by downloading samples with associated SRA accessions, resulting in 939 individual samples. Each of these was processed using *bronko* using a database with 2 representative strains from each Nextstrain [44] clade of SARS-CoV-2 (the first and last by sampling date, filtered for complete genomes with no ambiguous bases). This resulted in 100 total references from many different clades and timepoints throughout the pandemic. After quality filtering for breadth and depth of coverage, we were left with 372 samples. Of these, only 58 individuals contained more than 1 sample, reducing the final number of samples included in our analyses to 125.

Across these 125 samples, 853 total mutations were identified with respect to the selected reference genomes, comprising of 421 major variants (MAF ≥ 0.5) and 432 minor variants (0.01 ≤ MAF *<* 0.5). Among them, 524 (150 major, 374 minor) were unique within an individual patient, while the remaining 329 were recurrent within individual patients (271 major variants across 130 individuals/positions combinations; 58 minor variants across 30 individuals/positions). This pattern supports prior observations [5], that major variants tend to persist in an individual whereas minor variants are typically more transient. Additionally, as seen in **Figure 4A/B**, both the number of major variants and number of minor variants remained reasonably stable across samples, except for individuals with longer infection periods (>100 days), suggesting that substantial diversification of SARS-Cov-2 does not occur in a typical infection period (<14 days).

**Fig. 4.**
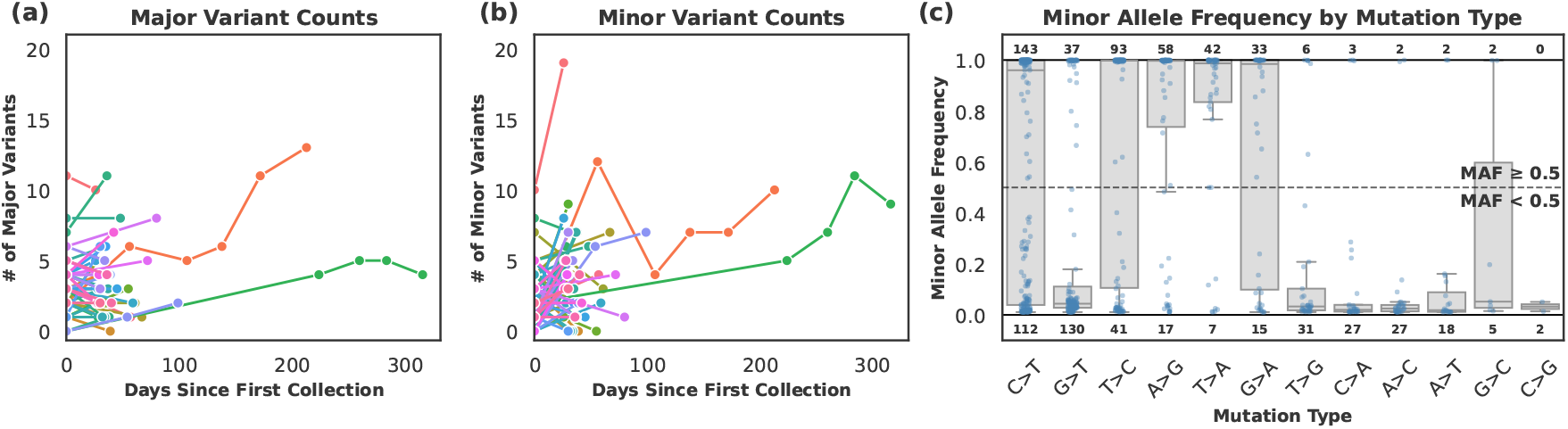
Analysis of chronically infected patients with SARS-CoV-2 **a**. Number of major variants for each individual across subsequent sampling periods **b** Number of minor variants for each individual across subsequent sampling periods **c** Frequency of major and minor mutations for each possible substitution. Mutations are sorted from left to right in order of the total number of variants. The number of major variants (>=50% MAF) is shown above the boxplot, while the sum of minor variants (<50% MAF) is shown below

We next examined within-host temporal shifts between variant classes (a minor variant transitioning into a major variant, or vice versa) across timepoints within an individual. Across all mutations and patients, we found 8 such transitions among four individuals. One individual, shown in orange in **Figure 4A/B** was responsible for four of these, of which two developed into a major variant from below 20% MAF. Three other patients contributed two, one, and one additional minor to major variant transitions. Of these, none developed from below 20% mutation rate, with all four initially being identified between 20 and 50% MAF. While it is difficult to make significant conclusions from such a small sample, these cases reiterate the value of tracking minor variants that could develop into major variants, even despite their generally transient nature.

Finally, we characterized the spectrum of allele frequencies across mutation types for both major and minor mutations (see **Figure 4C**). Again, in alignment with many previous studies [45, 46], C>T, G>T, and T>C, and A>G transitions dominated. Although this example is only a small sample set in chronically infected individuals, we observed that proportionally, 5 mutations (C>G, T>C, A>G, T>A, and G<A) had more major variants than minor, suggesting that these mutations may be more likely to establish themselves in an individual. On the other hand the remainder, including the 2nd most common G>T mutation, less commonly established as major variants, suggesting are rarely advantageous for the virus. The other potential explanation is that these variants are more likely to be artifacts of the sequencing and PCR process.

## 4 Discussion

In this work we introduced *bronko*, an alignment-free framework for identifying viral genome variation directly from sequencing reads. By constructing pileups from k-mers with small edit distance to the reference instead of relying on read alignment, *bronko* bypasses the computational bottlenecks inherent to traditional read-alignment based pipelines. Across simulated viral sequencing datasets, *bronko* achieved comparable precision in identifying minor variants to established variant calling tools such as LoFreq and iVar, while operating several orders of magnitude faster. *bronko* also was able to reconstruct alignments across larger sample sets that compared favorably to classical consensus-generation and core-genome alignment, with challenges only arising when the divergence between genomes was very high. In practice, these limitations should be mitigated by *bronko*’s flexible reference database, which allows users to leverage existing sets of viral reference sequences that are related to the given sample population. This reduces the cases of high divergence between the query and reference and minimizes the practical impact of indels, which bronko does not explicitly model and thus is unable to call. This framework also allows each reference to be queried individually against each sample, enabling rapid reference selection and SNP variant calling across large-scale studies of diverse lineages.

*bronko*’s use of a (1,2) LSB function allows it to identify mutations as long as there are not 3 variants within a k-length region. However, even in this case, both flanking variants would be identified, and only the middle of the three variants would be missed. This is a sensitivity improvement over split-k-mer based methods such as kSNP and SKA2, which at their core fail to detect any variants occurring within 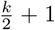 bases of another variant. While these methods are generally well-suited to bacterial genomes, where larger genome sizes, higher divergence, and more dispersed variation make missed sites less consequential, viral genomes demand greater sensitivity in many cases, particularly if divergence increases towards and beyond 1%. In practice, however, several parameters can influence this theoretical sensitivity. Most obviously, the k-mer size (default 21) directly influences the sensitivity and specificity, and is configurable by the user. To mitigate sequencing artifacts near the ends of reads, which can often present at low frequency, *bronko* by default also fixes bases at the end of k-mers such that they cannot be used to detect variants. This approach reduces the number of spurious variants that can appear, but in turn also reduces sensitivity in variant-dense regions. This tradeoff aligns with our general principle in designing *bronko* for precision over sensitivity, particularly for minor variants. Nonetheless, even with the fixed bases at the end of reads, we are still able to identify variants as long as there is no variant within the number of fixed bases plus the number of expected supporting k-mers (by default 2+2), still outperforming split-k-mer based methods. For users interested in maximum sensitivity when identifying major variants, these parameters can simply be set to zero to achieve the maximum theoretical sensitivity described above.

Additionally, as proposed in [31, 47], several alternative LSB functions exist beyond the (1,2) scheme implemented in *bronko*. These variants of LSB functions could, in principle, improve sensitivity in capturing clustered regions of local diversity. However, the strength of the (1,2) function lies in its simplicity, with each bucket directly corresponding to a potential SNP. In contrast, more complex LSB functions (ex. (2,3)) introduce additional complexity, such as more than k buckets per k-mer, which complicates the mapping protocols. Additionally, allowing for more distant matches increases the potential for unintentional hits, which may necessitate a larger k-mer size or other algorithmic adjustments. Exploring these alternative functions, therefore, presents an interesting direction for future work, although it remains to be seen whether the increased function complexity would yield meaningful gains in performance. Alternatively, expanding *bronko*’s existing framework to perform local assembly over variant dense regions provides a separate opportunity for improvement, and offers the ability to identify novel indels. We also aim to scale *bronko*’s framework to larger genomes. At present, the runtime and memory costs of enumerating each k-mer *k* times makes *bronko* more well-suited for viral genomes and other small targets, however ongoing optimization could further extend efficient support to larger bacterial genomes and other larger targets. Finally, while we showed our variant calling approach performs well, there are many other potential avenues for improvement, including integrating explicit models on sequencing and PCR errors, or utilizing existing datasets to classify artifacts from real signal [48].

## 5 Acknowledgments

R.D. is supported by a training fellowship from the Gulf Coast Consortia, on the NLM Training Program in Biomedical Informatics & Data Science (T15LM007093). M.J.T is supported for this effort by NIH grant 1U54AG089335-01. T.J.T is supported by NIH grant P01-AI152999 and NSF awards IIS-2239114 and EF-2126387. This work was also supported in part by the Texas Medical Center Genomic Center for Infectious Diseases (TMC-GCID) through NIH NIAID grant 1U19AI144297. We would also like to thank Dr. Michael Nute for his valuable feedback on the manuscript. Lastly, the authors would like to thank the anonymous RECOMB reviewers for their helpful comments and feedback

