## Supplementary Materials for "bronko: ultrafast, alignment-free detection of viral genome variation"

### 1 Appendix

#### 1.1 GENOMICON-Seq Parameters

We used GENOMICON-Seq v1.0 to simulate amplicon sequencing data. Ten HPV samples were generated using the default configuration, although mutation targets were removed. For each sample, we produced a single replicate and 1 million paired-end 250 bp reads under a NovaSeq error model (seed = 893). For performance evaluation, we generated additional samples with identical settings but varied sequencing depth: 100k, 500k, 5M, and 10M reads.

#### 1.2 Benchmarking *bronko build*

All results in Figure 3 are shown using a single reference. However, *bronko* can be utilized in many different ways, including varying genome sizes and incorporating multiple references of the same species. In the following section, we report on the effects of different configurations on runtime and memory usage. All commands were run using 10 threads.

First, we investigate the impact of genome size on bronko's memory usage and runtime. We generated random genomes of lengths 500, 1000, 2500, 5000, 10000, 25000, 50000, 100000, 250000, 500000, 1000000, and 2500000, and tested the effect on *bronko*'s build command. As seen in **Supp Fig. 1a/b/c**, using a single genome reference does not have a substantial effect on the runtime, memory usage, or index size within the range of typically-sized viral genomes. Even for a 1Mbp genome, the build step only took 31 seconds and 2.5GB, producing an index of 397MB. For most viral genomes of interest (SARS-CoV-2, Influenza, etc), genome lengths are closer to the range of 5-50kbp, and therefore do not exhibit a substantial effect on runtime, memory usage, or index size. We also investigated the effect a larger genome size (and therefore index) would have on the downstream variant calling. We tested *bronko* call on each index built from the genomes using simulated 250bp reads up to specified depths. As seen in **Supp Fig. 1d/e**, the call step was also reasonably unimpacted by the size of the genome, with only large genomes with high depth exhibiting substantial affect on runtime. Again, for most typical viral genome lengths, runtime remained below 10-15 seconds and memory usage remained around 200MB.

*bronko* also enables users to build a database of multiple reference sequences (for example lineages of interest from a specific species). Therefore, we also investigated the impact of the number of genomes on bronko's memory usage and runtime. In this example, we downloaded 100 SARS-CoV-2 genomes, and built bronko databases with increasing subsets of those genomes. As seen in **Supp Fig. 2a/b/c**, the runtime, memory usage, and index size all scale linearly with the number of genomes. When using all 100 genomes, the build step still only took 20 seconds and 6GB, producing an index of 449MB. Similar to before, we also investigated the effect of more references on downstream variant calling. We tested *bronko* call on each index againt simulated SARS-CoV-2 sequencing data of increasing depth. As seen in **Supp Fig. 2d/e**, the call step was also reasonably unimpacted by the number of genomes. Even when using all 100 genomes against 30000x data, the runtime remained near 15 seconds and memory usage was only 2GB.

#### 1.3 Simulated data for HIV multiple sequence alignment experiments

Reads were generated at 15,000x depth for each set using the same procedure as for the SARS-CoV-2 experiment. Test set S1 represents the most challenging case for *bronko*, with highly divergent and mutation-dense HIV sequences. Parsnp2 identified between 107 and 351 mutations (median 222) across the 41 samples, representing near 3% divergence from the reference on average. In this setting, *bronko* still identified 9041/11226 (88.4%) of the variants identified by Parsnp2, and detected 11 additional variants. Of the 1175 variants missed by *bronko*, 826 (70.3%) were in regions with 3 SNPs within k bases (3-in-k). Altogether, this data suggests that even in cases where newly sequenced viruses are very divergent from known references, *bronko* still recovers the majority of major variants aside from certain mutation-dense regions. Test set S2 represents a moderate divergence scenario, with 57-134 (median 101) variants per sample, representing around 1% divergence. Here, *bronko* was able to identify 3754/3888 variants (96.5%), where 88 of the 134 (65.6%) missed variants was due to the 3-in-k limitation. Finally, test set S3 represents the simplest and most representative case, with between 6-28 (mean 18) mutations per sample, representing an 0.2% average divergence. Here, *bronko* had perfect alignment with Parsnp2, identifying 664/664 variants. Although *bronko* exhibits slight dropout at high divergence, in outbreak settings, most viral sequencing data are closely related (<1%) to available references, and *bronko* is designed to query many references in parallel to mitigate this issue. We

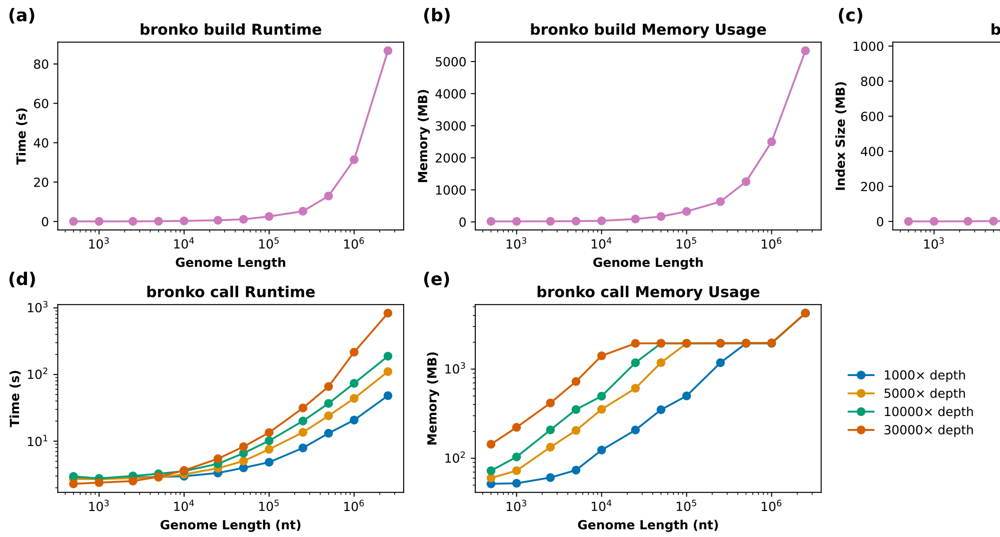


**Supp Fig. 1.** Benchmarking *bronko* across different genome sizes of randomly simulated sequences **a/b**. Runtime and memory usage of *bronko* build command across different genome sizes **c** *bronko* index size across different genome sizes **d/e** Runtime and memory usage of *bronko* call command size across different genome sizes and sequencing depths

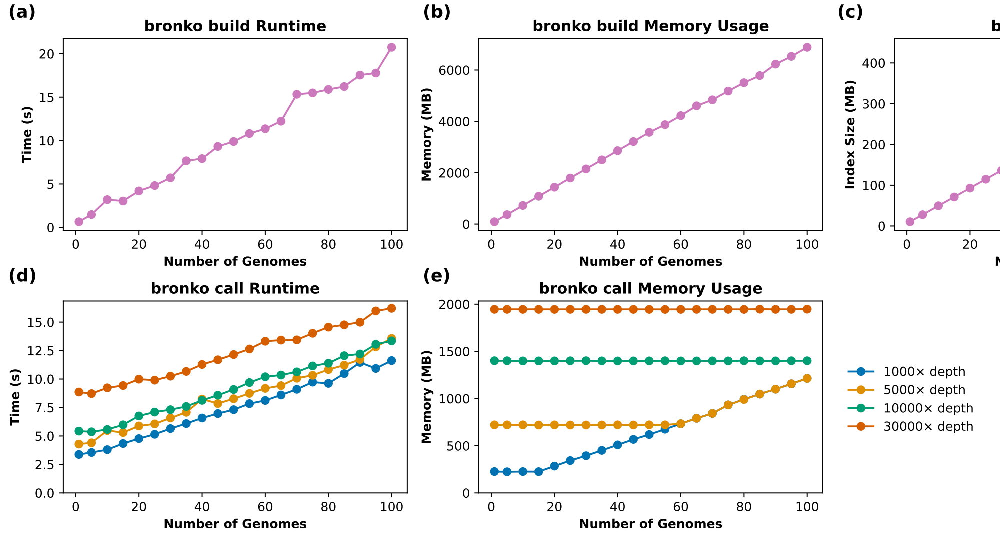


**Supp Fig. 2.** Benchmarking *bronko* with increasing numbers of genomes in the .bkdb database **a/b**. Runtime and memory usage of *bronko* build command with more genomes **c** *bronko* index size with more genomes **d/e** Runtime and memory usage of *bronko* call command size with more genomes and different sequencing depths

also quanitified the total number of variants across all datasets. This included the number occurring in a 3-in-k region (theoretical limit of *bronko*), as well as the number occurring in regions with 2 variants within $(\frac{k}{2}+1)$ bases, which represents the theoretical limitation of split-kmer based approaches (**Suppplementary Table 1**). Across all three HIV test sets, *bronko* ran over 5x faster, although utilized 2x more memory. However, again, this comparison does not account for the fact that the alignment pipeline does not detect minor variants, which would substantially increase runtime, as shown in Figure 3.

**Supplementary Table 1.** Simulated data SNP presence and density overview (k=21)

| **Dataset (#)** | **Total Parsnp2 SNPs** | **Median SNPs per sample** | **SNP Range (min-max)** | **2-in-($\frac{k}{2}+1$) SNPs** | **3-in-k SNPs** |
|---|---|---|---|---|---|
| CoV (n=543) | 3647 | 6 | 2-14 | 4 | 0 |
| HIV (S1) (n=41) | 10226 | 222 | 107-351 | 4289 | 2406 |
| HIV (S2) (n=39) | 3888 | 101 | 57-134 | 983 | 286 |
| HIV (S3) (n=37) | 664 | 18 | 6-28 | 12 | 0 |

##### 1.4 Accessions for Longitudinal SARS-Cov-2 Data

All data for the longitudinal SARS-CoV-2 data is available under the BioProject PRJEB37886. However, the specific accessions selected in this study are in the bronko-test repository.
